# Navigating Neurological Re-emergence in Feline Infectious Peritonitis: Challenges and Insights from GS-441524 and Remdesivir Treatment

**DOI:** 10.1101/2025.04.04.646959

**Authors:** Celia C de Witt Curtius, Maxime Rodary, Regina Hofmann-Lehmann, Andrea M Spiri, Marina L Meli, Aline Crespo Bouzon, Jennifer Wenk, Ilaria Cerchiaro, Benita Pineroli, Simon A Pot, Katrin Beckmann, Tatjana Chan, Manuela Wieser, Stefan Unterer, Sandra Felten, Solène M Meunier

## Abstract

**Case summary:** A six-month-old male British Longhair cat presented with acute neurological signs, ocular changes, massive ascites and laboratory parameters consistent with feline infectious peritonitis (FIP). Systemic and neurological signs fully resolved with initial treatment (GS-441524 [BOVA UK, 15 mg/kg PO q24h, 42 days], levetiracetam [20 mg/kg q8h] and prednisolone [1 mg/kg q24h until Day 21 = D-21]). Lethargy and fever reappeared seventeen days after treatment. Four days later, severe multifocal neurological signs reemerged. High-field MRI revealed multifocal intra-axial and intramedullary lesions in the brainstem and cervical spinal cord, severe meningitis and generalised mild ventriculomegaly. Feline coronavirus (FCoV) RNA was detected in the cerebrospinal fluid by RT-qPCR. Abdominal effusion was absent. Serum alpha-1-acid glycoprotein (AGP) was again elevated. FIP re-emergence was suspected, and antiviral treatment was resumed. After one day of GS-441524 treatment (15 mg/kg PO q24h), severe hypoventilation developed, requiring intubation and mechanical ventilation for 1.5 days. Treatment was switched to remdesivir (Veklury, Gilead, 16.7 mg/kg IV q24h) for four days. Oral GS-441524 was then reintroduced (10 mg/kg q12h) and continued until D-84. Treatment resulted in partial recovery with moderate ataxia and reduced left-sided menace response remaining 181 days after starting the second treatment.

**Relevance and novel information:** This case illustrates the complexity of diagnosing and treating re-emerging FIP-associated neurological signs. AGP monitoring offers a promising noninvasive approach for early detection of relapse. By adapting short- and long-term antiviral treatment and providing intensive care, excellent long-term outcomes can be obtained for cats with severe relapsing FIP-related neurological signs.

## 1. Introduction

Although new effective oral antiviral drugs often lead to excellent short-term recovery and long-term remission in cats with FIP, the recurrence of clinical signs is possible.^1-5^ Cats with FIP-related neurological manifestations are considered harder to treat and have a higher risk of recurrence due to the presumed limited drug penetration across the blood-brain barrier.^6,7^ Therefore, careful monitoring during and after antiviral treatment is crucial. MRI and cerebrospinal fluid (CSF) analysis have proven useful but are associated with limited availability, high costs and the need for general anaesthesia.^6^ Noninvasive monitoring of serum alpha-1-acid glycoprotein (AGP) and serum amyloid A (SAA) levels could benefit cats with FIP-related neurological manifestations, as with those with cavitary effusions.^8^ Evidence is lacking regarding the optimal treatment for recurrent FIP-associated neurological signs. We describe the follow-up and successful treatment of a cat with re-emergent FIP-associated neurological signs despite initial successful oral antiviral treatment with GS-441524.

## 2. Case Description

### 2.1. Signalment and history

A six-month-old intact male British Longhair cat, kept indoors with a partner, initially presented with acute-onset neurological signs, ocular changes and abdominal distension. On admission, the cat showed lethargy, ventroflexion, severe bilateral uveitis, active chorioretinitis, mild ataxia and a single self-limiting generalised tonic-clonic seizure (Table 1). Complete blood count (CBC) and serum chemistry, including elevated AGP (4966 μg/ml) and SAA (80.6 mg/l), suggested FIP (Table 2, Figure 1). Abdominal ultrasound revealed generalised lymphadenomegaly and massive corpuscular ascites. Abdominocentesis yielded protein-rich transudate with a positive Rivalta test and AGP of 3650 μg/ml. High positive FCoV reverse transcription quantitative PCR (FCoV RT-qPCR) confirmed FIP, with viral loads of 3.0 × 10^8^ copies/ml in ascites and 5.9 × 10^5^ copies/ml in blood.^9^ Central nervous system (CNS) imaging and CSF analysis were not performed. Oral antiviral treatment was initiated (GS-441524, BOVA UK; 15 mg/kg q24h, 42 days). Supportive care included fluids, maropitant, ondansetron, mirtazapine, prednisolone (1 mg/kg q24h until D-21, tapered), long-term levetiracetam (20 mg/kg q8h), and ophthalmological treatment. This led to full systemic and neurological resolution, though permanent, inactive chorioretinal and iris-lens lesions remained (Table 1). By D-42, all blood parameters had normalized and FCoV RT-qPCR from blood detected no FCoV viral RNA, supporting the decision to discontinue treatment (Figure 1).

**Table 1.** Clinical and neurological signs at initial presentation (D-1), at the end of the first treatment (D-42) and at the time of diagnosis of FIP re-emergence (D-66/1)

| Parameter | D-1 (initial presentation) | D-42 (end of first treatment) | D-66/1 (re-emergent FIP) |
| --- | --- | --- | --- |
| Body Condition Score | 2/9 | 3/9 | 4/9 |
| Mental status | Lethargic, responsive | Alert, responsive | Obtunded |
| Posture | Ventroflexion | Normal posture | Intermittent left-sided head tilt |
| Gait | Slight ataxia | Normal gait | Nonambulatory tetraparesis |
| Proprioception/<br>Hopping | Prompt (all four limbs) | Prompt (all four limbs) | Reduced (all four limbs) |
| Cranial nerves | Multiple deficits | Normal | Absent menace response, oculocephalic and nasocortical reaction |
| Seizures | Tonic-clonic seizure | No seizures | No seizures |
| Ascites | Severe | Mild (not procurable) | None |
| Ocular status | Uveitis, chorioretinitis | Healed uveitis, inactive chorioretinitis | Not performed |

D-1 = Start of the initial FIP treatment, D-42 = End of the initial FIP treatment, D-66/1 = Day of presentation at our hospital due to re-emerging FIP clinical signs

**Table 2.**
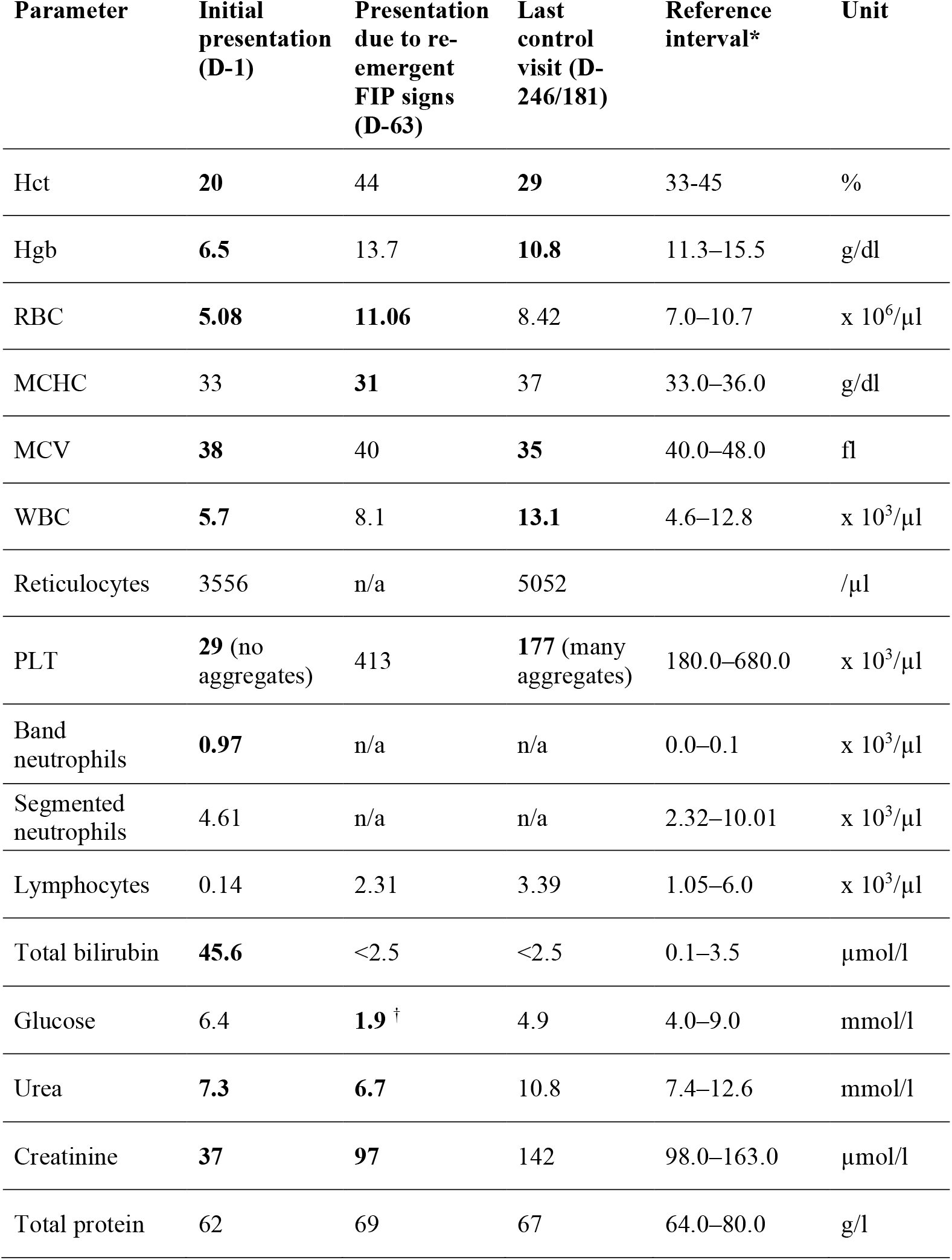
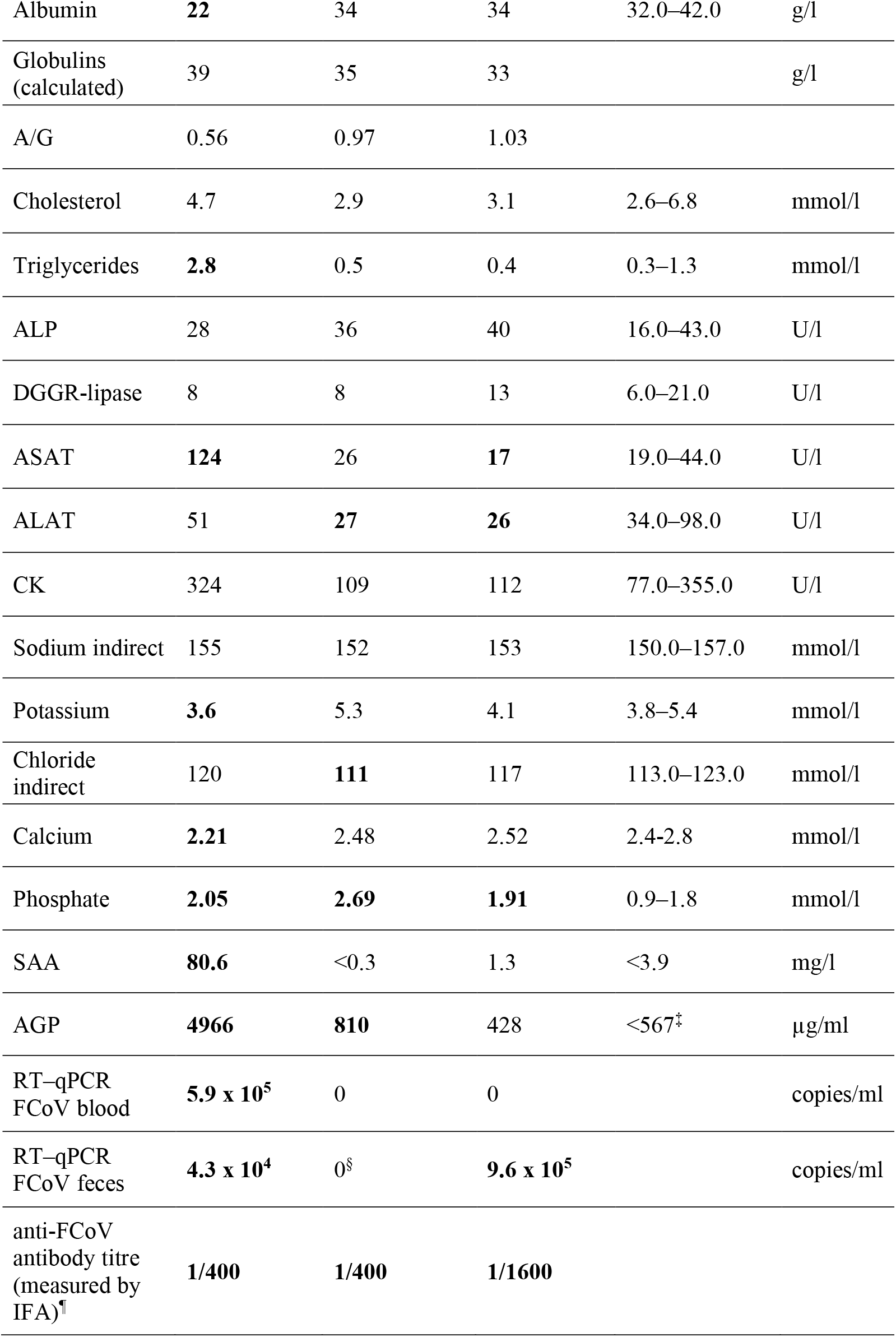
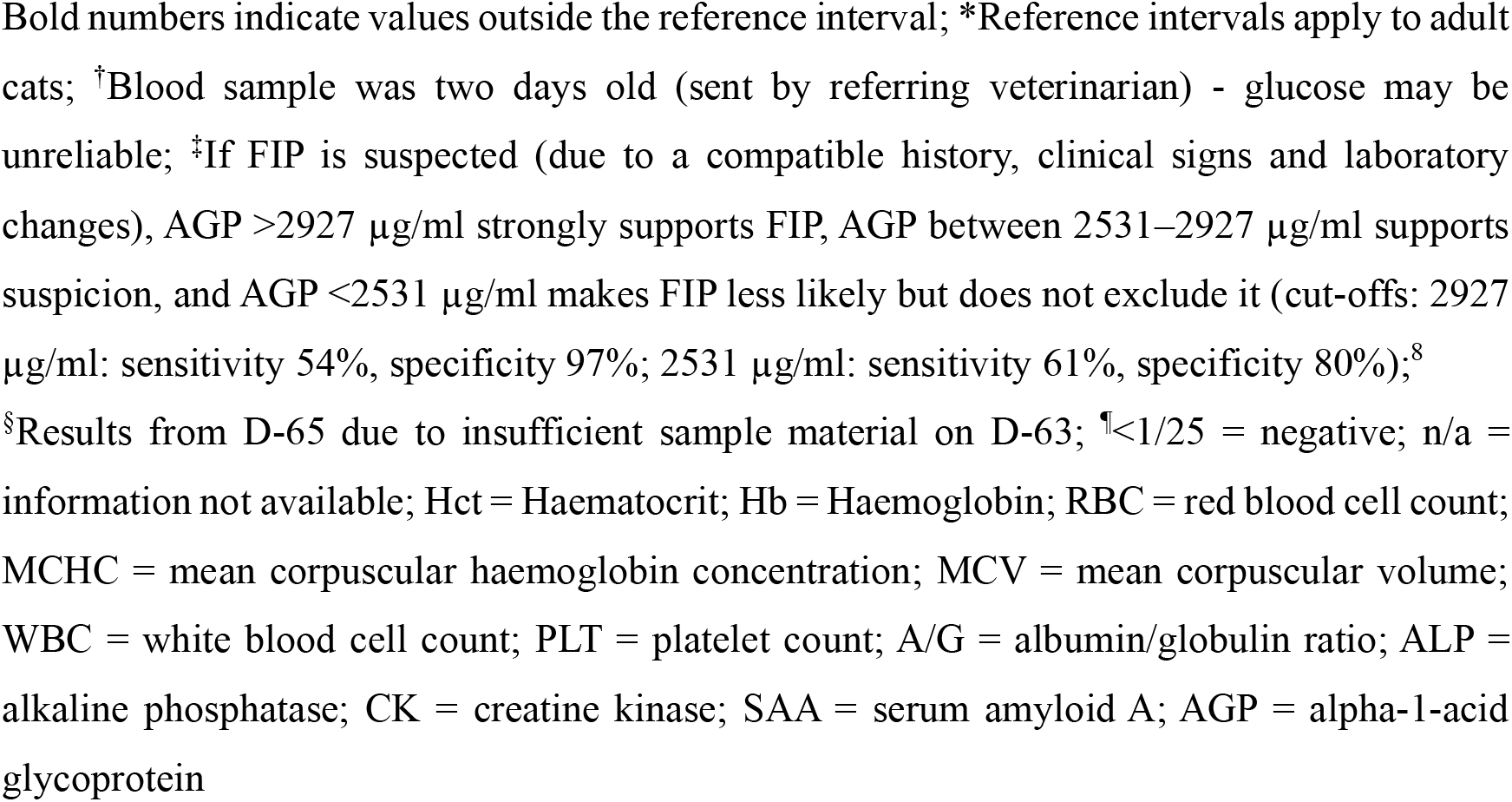
Comparison of blood work at the initial presentation (D-1 of first treatment), at the time of re-emergence of feline infectious peritonitis (FIP)-associated clinical signs (D-63) and at the last follow-up (D-246/181 of first/second treatment, respectively)

| Parameter | Initial presentation (D-1) | Presentation due to re-emergent FIP signs (D-63) | Last control visit (D-246/181) | Reference interval* | Unit |
| --- | --- | --- | --- | --- | --- |
| Hct | **20** | 44 | **29** | 33–45 | % |
| Hgb | **6.5** | 13.7 | **10.8** | 11.3–15.5 | g/dl |
| RBC | **5.08** | **11.06** | 8.42 | 7.0–10.7 | x 10<sup>6</sup>/μl |
| MCHC | 33 | **31** | 37 | 33.0–36.0 | g/dl |
| MCV | **38** | 40 | **35** | 40.0–48.0 | fl |
| WBC | **5.7** | 8.1 | **13.1** | 4.6–12.8 | x 10<sup>3</sup>/μl |
| Reticulocytes | 3556 | n/a | 5052 |  | /μl |
| PLT | **29** (no aggregates) | 413 | **177** (many aggregates) | 180.0–680.0 | x 10<sup>3</sup>/μl |
| Band neutrophils | **0.97** | n/a | n/a | 0.0–0.1 | x 10<sup>3</sup>/μl |
| Segmented neutrophils | 4.61 | n/a | n/a | 2.32–10.01 | x 10<sup>3</sup>/μl |
| Lymphocytes | 0.14 | 2.31 | 3.39 | 1.05–6.0 | x 10<sup>3</sup>/μl |
| Total bilirubin | **45.6** | <2.5 | <2.5 | 0.1–3.5 | μmol/l |
| Glucose | 6.4 | **1.9** † | 4.9 | 4.0–9.0 | mmol/l |
| Urea | **7.3** | **6.7** | 10.8 | 7.4–12.6 | mmol/l |
| Creatinine | **37** | **97** | 142 | 98.0–163.0 | μmol/l |
| Total protein | 62 | 69 | 67 | 64.0–80.0 | g/l |
| Albumin | **22** | 34 | 34 | 32.0–42.0 | g/l |
| Globulins<br>(calculated) | 39 | 35 | 33 |  | g/l |
| A/G | 0.56 | 0.97 | 1.03 |  |  |
| Cholesterol | 4.7 | 2.9 | 3.1 | 2.6–6.8 | mmol/l |
| Triglycerides | **2.8** | 0.5 | 0.4 | 0.3–1.3 | mmol/l |
| ALP | 28 | 36 | 40 | 16.0–43.0 | U/l |
| DGGR-lipase | 8 | 8 | 13 | 6.0–21.0 | U/l |
| ASAT | **124** | 26 | **17** | 19.0–44.0 | U/l |
| ALAT | 51 | **27** | **26** | 34.0–98.0 | U/l |
| CK | 324 | 109 | 112 | 77.0–355.0 | U/l |
| Sodium indirect | 155 | 152 | 153 | 150.0–157.0 | mmol/l |
| Potassium | **3.6** | 5.3 | 4.1 | 3.8–5.4 | mmol/l |
| Chloride<br>indirect | 120 | **111** | 117 | 113.0–123.0 | mmol/l |
| Calcium | **2.21** | 2.48 | 2.52 | 2.4–2.8 | mmol/l |
| Phosphate | **2.05** | **2.69** | **1.91** | 0.9–1.8 | mmol/l |
| SAA | **80.6** | <0.3 | 1.3 | <3.9 | mg/l |
| AGP | **4966** | **810** | 428 | <567<sup>‡</sup> | µg/ml |
| RT–qPCR<br>FCoV blood | **5.9 x 10<sup>5</sup>** | 0 | 0 |  | copies/ml |
| RT–qPCR<br>FCoV feces | **4.3 x 10<sup>4</sup>** | 0<sup>§</sup> | **9.6 x 10<sup>5</sup>** |  | copies/ml |
| anti-FCoV<br>antibody titre<br>(measured by<br>IFA)<sup>¶</sup> | **1/400** | **1/400** | **1/1600** |  |  |

Bold numbers indicate values outside the reference interval; \*Reference intervals apply to adult cats; †Blood sample was two days old (sent by referring veterinarian) - glucose may be unreliable; ‡If FIP is suspected (due to a compatible history, clinical signs and laboratory changes), AGP >2927 µg/ml strongly supports FIP, AGP between 2531–2927 µg/ml supports suspicion, and AGP <2531 µg/ml makes FIP less likely but does not exclude it (cut-offs: 2927 µg/ml: sensitivity 54%, specificity 97%; 2531 µg/ml: sensitivity 61%, specificity 80%);<sup>8</sup>
§Results from D-65 due to insufficient sample material on D-63; ¶<1/25 = negative; n/a = information not available; Hct = Haematocrit; Hb = Haemoglobin; RBC = red blood cell count; MCHC = mean corpuscular haemoglobin concentration; MCV = mean corpuscular volume; WBC = white blood cell count; PLT = platelet count; A/G = albumin/globulin ratio; ALP = alkaline phosphatase; CK = creatine kinase; SAA = serum amyloid A; AGP = alpha-1-acid glycoprotein

**Figure 1.**
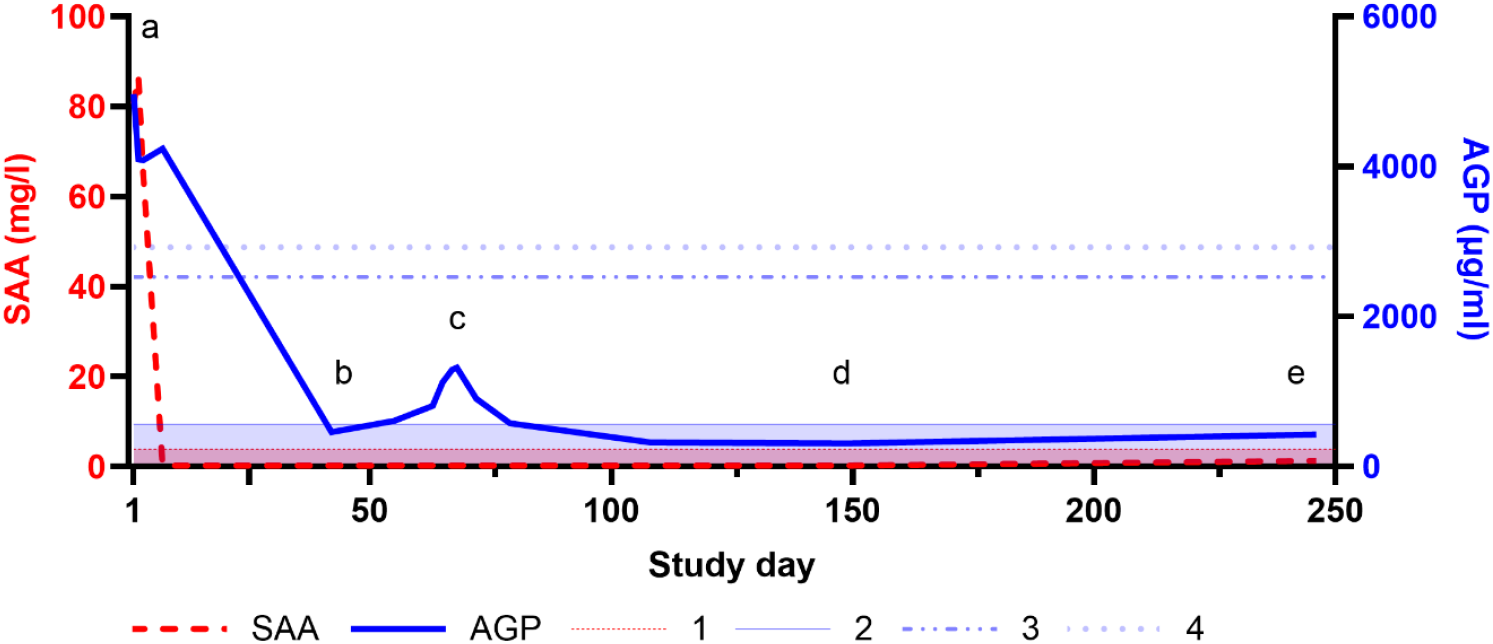
Time course of serum amyloid A (SAA) (thick dashed line, on the left side) and serum alpha-1-acid glycoprotein (AGP) (thick solid line, on the right side) concentrations. 1 = Reference interval SAA: <3.9 mg/l; 2 = Reference interval AGP: <567 µg/ml; 3 = AGP between 2531–2927 µg/ml supports suspicion of FIP in a cat with compatible clinical signs (sensitivity 61%, specificity 80%); 4 = AGP >2927 µg/ml strongly supports FIP in a cat with compatible clinical signs (sensitivity 54%, specificity 97%);^8^ (a) Initial presentation (D-1); (b) End of first treatment (D-42); (c) Re-emergent FIP signs (D-63); (d) End of second treatment (D-149/84); (e) Last recheck (D-246/181)

### 2.2. Re-emergent FIP signs

Seventeen days posttreatment (D-59), lethargy and elevated temperature were reported. The cat was presented 21 days posttreatment (D-63) to the referring veterinarian with rapidly progressing ataxia. Neurological examination on D-66 at our hospital revealed severe deficits consistent with multifocal neuroanatomical localisation including the brainstem (Table 1). Owing to poor clinical conditions, no ophthalmologic examination was performed.

### 2.3. Diagnostic imaging and laboratory findings at re-emergent FIP

On D-63, blood analysis revealed only elevated AGP (810 μg/ml, Table 2), with no effusion present. High-field MRI of the brain and cervical spine was performed on D-66/1 (days of initiation of the first/second treatment, respectively; beginning from now, time points will be given in this format). Intramedullary lesions extending from the medulla oblongata to C5, spinal cord swelling (Figure 2), moderate ventriculomegaly (Figure 3) and multifocal thickening of the meninges with strong contrast enhancement (Figure 4) were found. The swelling restricted access to the occipital cistern, necessitating a lumbar CSF puncture instead of an atlantooccipital approach. CSF analysis revealed moderate lymphocytic pleocytosis (341 nucleated cells/µl) and severely elevated total protein levels (18.7 g/l). No infectious organisms or atypical cells were observed. FCoV RT-qPCR was lowly positive (2.5 × 10^3^ copies/ml), strongly suggesting FIP relapse with meningoencephalomyelitis. Viral sequencing of the spike gene (99 bp) revealed the M1058L mutation associated with systemic spread and/or FIP^10,11^ in effusion, blood (D-1) and CSF (D-66/1) but not in feces (D-1 and D-246/181) (Figure 5). In addition to the M1058L mutation, early blood, effusion and fecal sequences (D-1) matched the late fecal sample (D-246/181) at the nucleotide level. In contrast, the CSF sequence (D-66/1) differed by 14/99 nucleotides from the effusion and blood sequences (D-1), with 13 synonymous and one nonsynonymous mutation. Thus, the CSF sequence (D-66/1) differed from the blood and effusion sequences (D-1) by only 1/32 amino acids.

**Figure 2.**
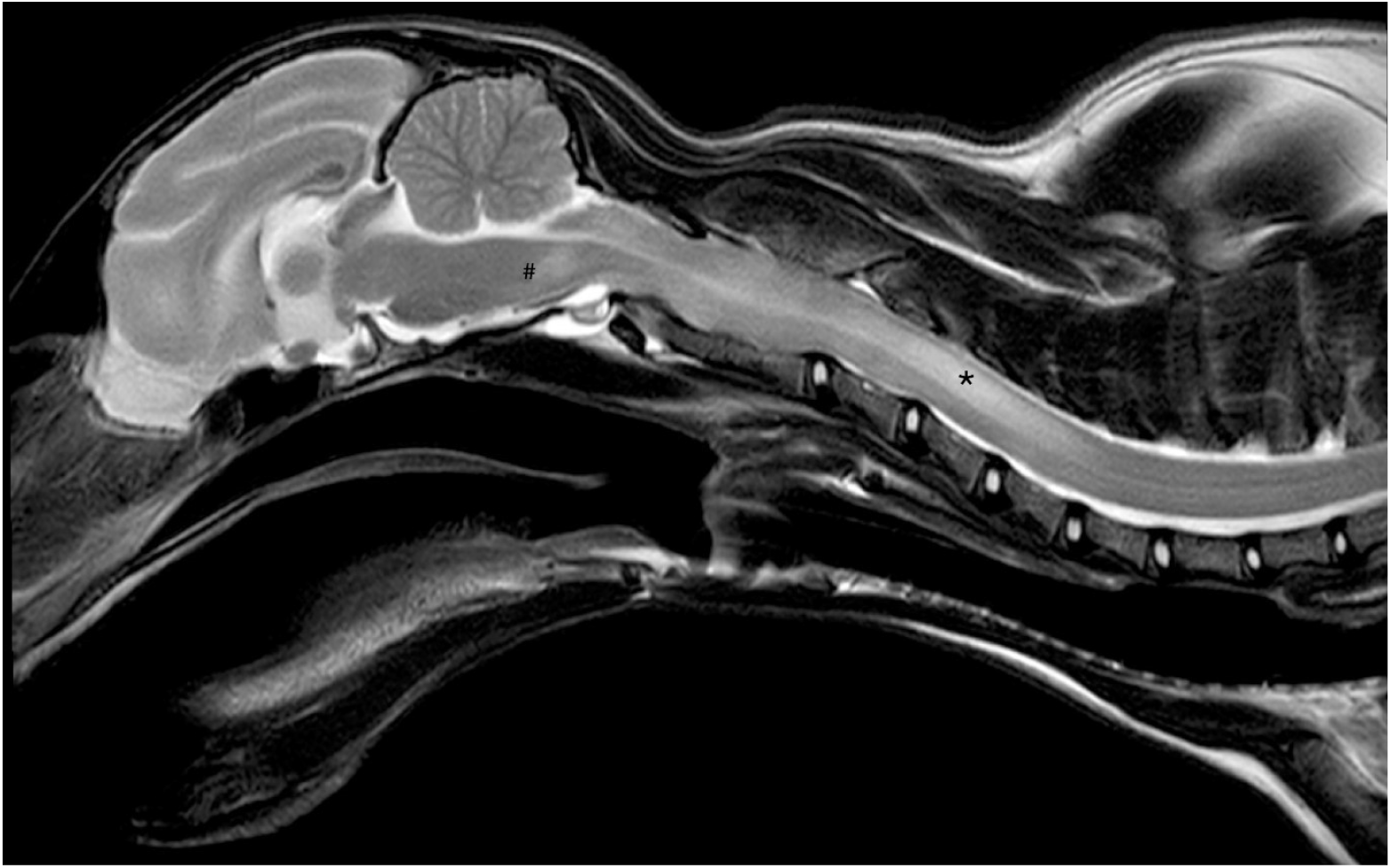
Sagittal T2-weighted (T2W) image of the brain and cervical spine. There was moderate spinal cord swelling and an extensive, asymmetrical, T2W and T1W hyperintense lesion in the mesencephalon (#) extending along the cervical spinal cord until C5. The lesion is most pronounced at the level of C4 (asterisk) © 2024 Prof. Patrick Kircher, Clinic for Diagnostic Imaging, Vetsuisse Faculty, UZH

**Figure 3.**
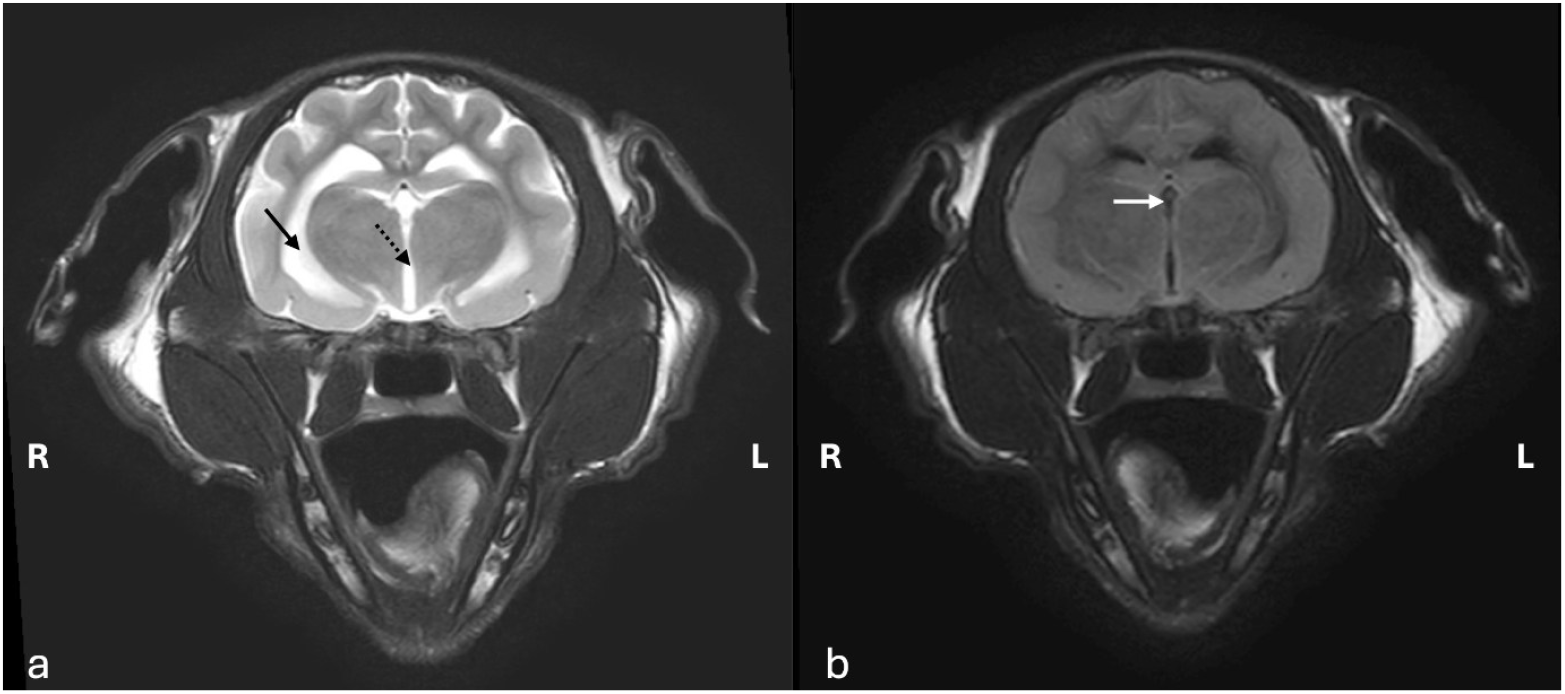
Transverse (a) T2-weighted (T2W) and (b) T2-fluid attenuation inversion recovery (FLAIR) images of the brain at the level of the thalamus and third ventricle. The ventricular system is moderately widened (arrow: lateral ventricle, dashed arrow: third ventricle) with incomplete CSF suppression in the FLAIR sequence (white arrow) © 2024 Prof. Patrick Kircher, Clinic for Diagnostic Imaging, Vetsuisse Faculty, UZH

**Figure 4.**
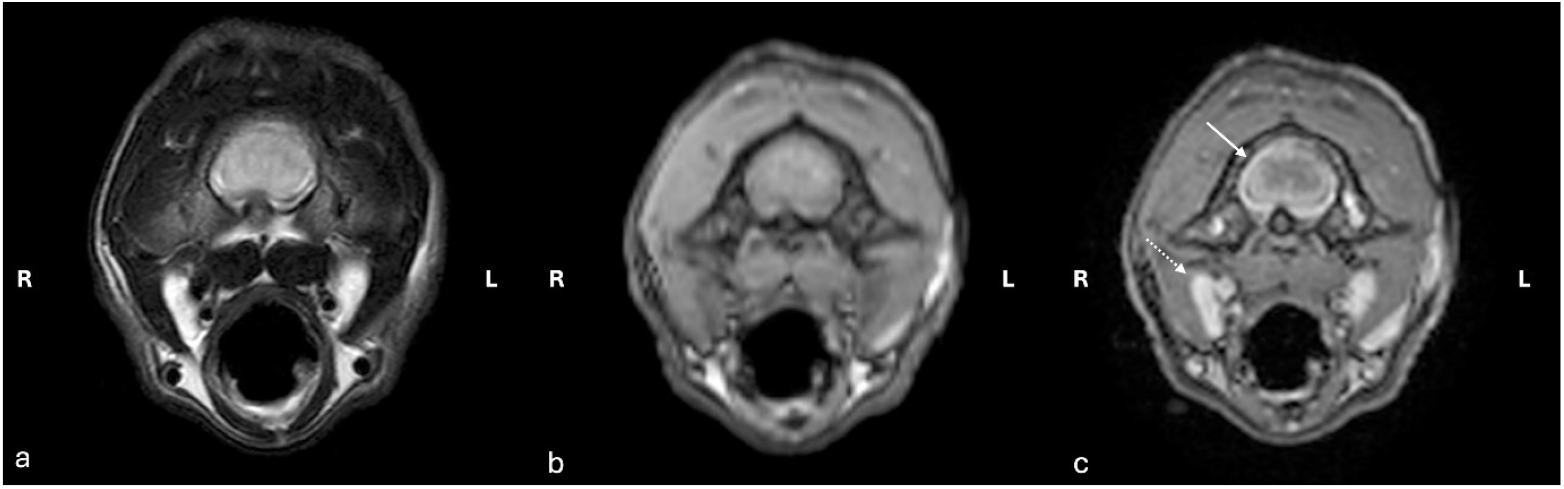
Transverse (a) T2-weighted (T2W), (b) T1-weighted (T1W) precontrast, and (c) T1-weighted (T1W C+) postcontrast images of the cervical spinal cord at the level of C1. The meninges are moderately thickened with strong contrast enhancement (white arrow), which is compatible with marked meningitis. Mild bilateral medial retropharyngeal lymphadenomegaly (dashed white arrow) © 2024 Prof. Patrick Kircher, Clinic for Diagnostic Imaging, Vetsuisse Faculty, UZH

**Figure 5.**
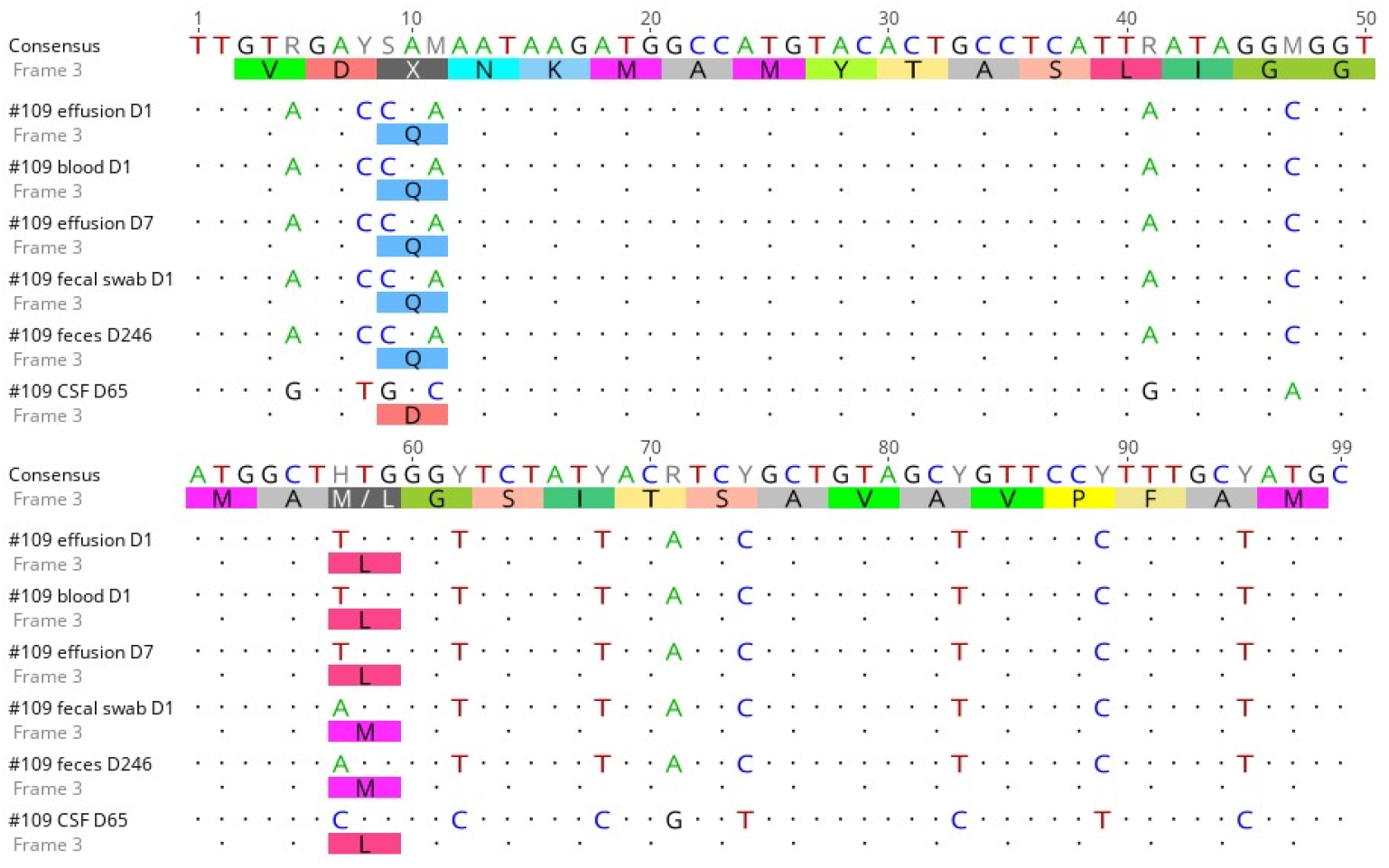
Nucleotide (upper line; Consensus) and translated amino acid sequences (bottom line; Frame 3) of the analysed FCoV strains. Dots depict the same nucleotide and amino acid sequences, whereas letters represent differences with respect to the consensus sequence. The sequences were edited and aligned via Clustal Omega,^13^ and images were created with Geneious Prime 2020.2.5

### 2.4. Treatment challenges of re-emergent FIP

Antiviral treatment was resumed (GS-441524, 15 mg/kg PO q24h) and mannitol was given once for suspected intracranial hypertension. On D-67/2, the cat’s mentation worsened, accompanied by bradypnea (<16 breaths per minute), shallow breathing, bradycardia (106 beats/minute) and hypothermia (35.1°C). Blood pressure remained normal. Owing to an absent swallowing reflex and severe hypercapnia (venous partial pressure of CO_2_ (pCO_2_) of 134.3 mmHg) secondary to hypoventilation, the cat was anesthetised, intubated and placed on positive pressure-controlled mechanical ventilation with a fraction of inspired oxygen (FiO_2_) of 30%. Total intravenous anaesthesia (TIVA) combined with propofol, dexmedetomidine and midazolam ensured deep sedation. Oral GS-441524 was replaced with intravenous remdesivir (Veklury, Gilead, 16.7 mg/kg q24h). After 22 hours, respiratory drive improved, allowing TIVA discontinuation and spontaneous breathing. Serial venous blood gas examinations at 4, 8 and 12 hours postextubation revealed pCO_2_ levels of 64.8, 79.3, and 76.8 mmHg, respectively. To prevent hypercapnia-associated hypoxemia, oxygen was supplied via an oxygen box (FiO_2_ 50%). The cat was tetraparetic with reduced mentation but had a normal respiratory rate and improved breathing. By D-69/4, the neurological status improved, allowing the transition to oral GS-441524 at an increased dose (10 mg/kg q12h). Oxygen supplementation was stopped when the haemoglobin saturation reached 100% under room air. Supportive medications included prednisolone (1 mg/kg IV q12h), levetiracetam, vitamin B1, ondansetron (until D-71/6), maropitant (until D-76/11) and mirtazapine (until D-67/2). Amoxicillin-clavulanic acid was initiated from D-67/2 until D-73/8 for suspected aspiration pneumonia.

### 2.5. Long-term follow-up

By D-77/12, the cat could stand and walk, although severe generalised ataxia persisted. The cat was discharged on D-79/14, but manual bladder expression remained necessary. Spontaneous micturition was first reported on D-83/18. Prednisolone was tapered and discontinued with D-100/35. By D-107/42, blood results were within reference intervals, and FCoV RT-qPCR from blood was negative. Owing to persistent neurological deficits, such as severe ataxia, slightly reduced tactile placing on the right side, a reduced menace response on the left side and a mildly reduced oculocephalic reflex (see video 1 in supplementary material), antiviral treatment was extended to 84 days. By D-149/84, owners reported the cat jumping onto windowsills and daily improvements in mobility and coordination (see video 2, 3 in supplementary material). The remaining reduced left-sided menace response, wide-based stance and moderate ataxia were attributed to remnant spinal cord damage and no longer associated with acute FIP. The second GS-441524 treatment was stopped. FCoV RT-qPCR from feces was weakly positive (D-149/84), contrasting expectation of complete FCoV absence after antiviral treatment.^12^ On D-246/181, only mild vestibular ataxia remained. Blood work was unremarkable, with mild deviations of questionable clinical relevance and a negative FCoV RT-qPCR result (Table 2).

## 3. Discussion

### 3.1. Relapse versus reinfection

When clinical signs of FIP re-emerge, distinguishing between relapse and reinfection may influence the therapeutic approach. A relapse can be defined as a recurrence due to incomplete viral clearance, possibly involving viral resistance. It occurs shortly after treatment, involves the same virus strain and may require higher dosages or longer treatment. Conversely, reinfection refers to a cat acquiring a new infection with a different or the same FCoV strain.^14^ Prior FCoV infection does not confer protective immunity, allowing reinfection despite high antibody levels.^15,16^ A reinfected cat may still respond to the initial treatment. In our case, although only a small portion of the FCoV virome was sequenced, the results suggest the presence of a similar virus in the CSF at D-66/1 (only one amino acid change in addition to synonymous mutations). The rapid reoccurrence of clinical signs and the absence of outdoor access or exposure to another FCoV support relapse. Identical fecal viral sequences from D-1 and D-246/181 indicate re-emergence but do not rule out reinfection by the same FCoV strain circulating in the household, as the partner cat may have been shedding FCoV but was not tested.

### 3.2. Risk factors and rate of re-emerging FIP

The incidence of re-emergent clinical signs during and after FIP treatment varies widely in the literature, ranging from 0 to 73.7% (Table 3).^1-5,17^ No comparison has been made between cats with and without initial FIP-related neurological manifestations. Known and suspected risk factors for FIP re-emergence can be grouped into cat-, virus-, clinical presentation- and treatment-related factors (Figure 6). These findings, along with varying follow-up lengths, probably explain the differences in the results of these studies. Among drug-related factors, the choice of antiviral agent may be relevant (Figure 6). In this case, lower GS-441524 concentrations in the CSF are thought to result from limited blood-brain barrier penetration, leading to the empirical recommendation of higher dosages for FIP-related neurological manifestations.^6,7,18^ By doing so, promising results were obtained in a recent study on unlicensed GS-441524-like treatment: 42.3% of the cats with neurological or ocular FIP-related manifestations responded positively, and only 4.7% died.^2^

**Table 3.** Reported rates of FIP re-emergence.

| Study | FIP re-emergence rate (% and number of cats with FIP-related neurological manifestations at re-emergence of clinical signs) | Study includes neurological cases | Number of cats (% and number of cats with FIP-related neurological manifestations at study inclusion, if applicable) | Treatment |
| --- | --- | --- | --- | --- |
| Ongoing bicenter study (UZH and LMU)<sup>∞</sup> | 1% (50%, 1/2) | yes | 202<sup>#</sup> (19.8%, 40/202) | GS-441524 & Remdesivir ** |
| Buchta et al., 2024<sup>∞</sup> | 2.5% (0%, 0/1) | no | 40 (n/a) | GS-441524** |
| Taylor et al., 2023 | 10.8% (51%, 17/33) | yes | 307 (20%, 62/306) | Remdesivir & GS-441524** |
| Coggins et al., 2023 | 16% (n/a) | no | 28 (n/a) | Remdesivir & GS-441524** |
| Jones et al., 2021 | 12.7% (n/a) | yes | 393 (26%, 103/393) | GS (unlicensed) |
| Krentz et al.,<br>2021 | 0%<br>(n/a) | no | 18 | GS-441524<br>(Xraphconn) |
| Yin et al.,<br>2021 | 3,3%<br>(n/a) | yes | 30<br>(n/a) | GS-441524 &<br>GC 376 |
| Pedersen et al.,<br>2017 | 73.7%<br>(n/a) | no | 19 | GC 376 |

<sup>∞</sup>unpublished data; <sup>#</sup>status as of 10/2024; <sup>\*\*</sup>(BOVA UK); n/a: information not available

**Figure 6.**
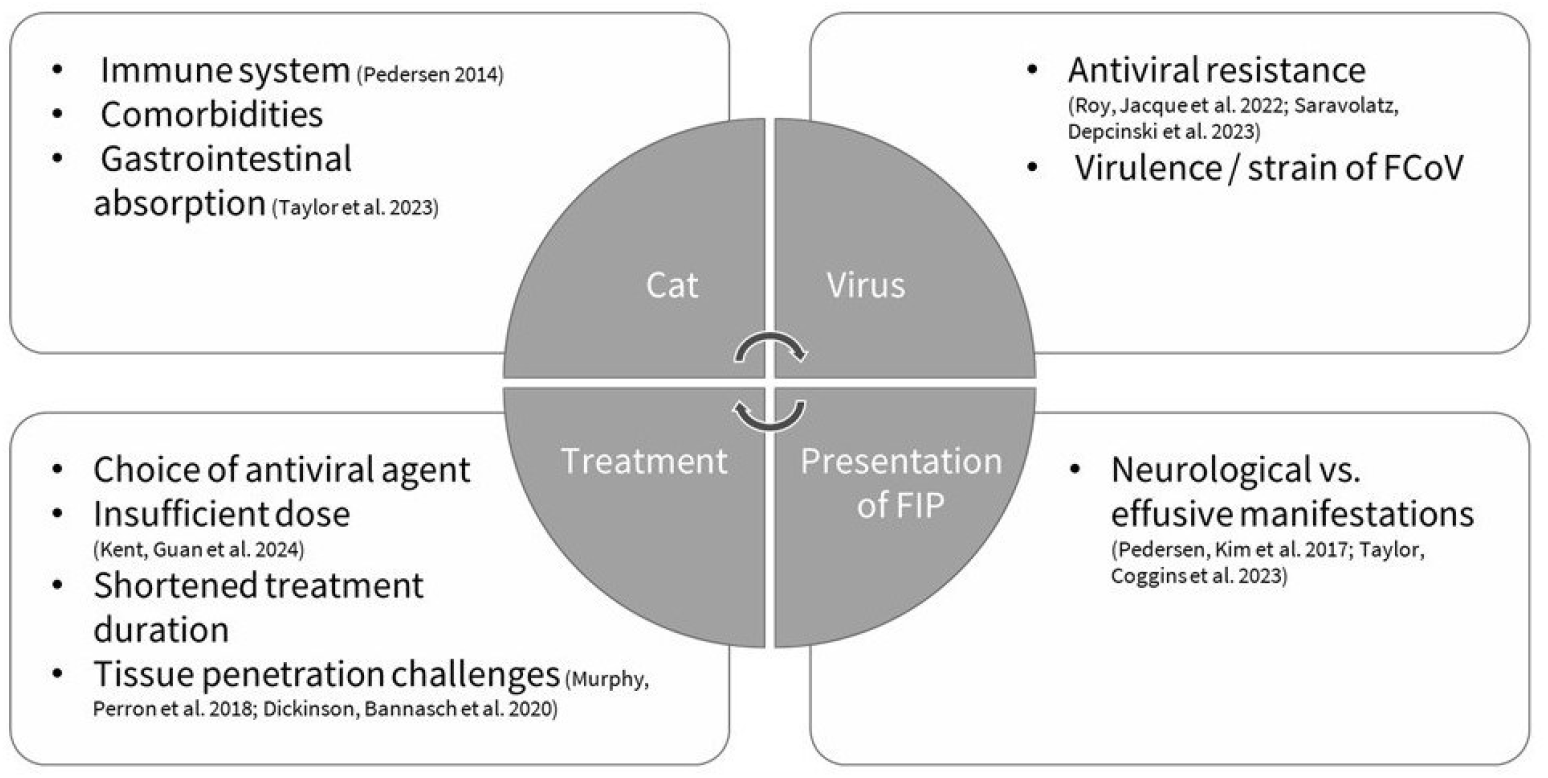
Known and suspected risk factors for FIP re-emergence

### 3.3. AGP, an early noninvasive monitoring biomarker

AGP is a moderate acute-phase protein that plays a critical role in the inflammatory response, with peak levels occurring approximately 2–3 days after activation.^19^ It has shown promise for monitoring treatment response and predicting FIP re-emergence.^8^ Retrospective analysis of AGP levels, in this case, revealed a gradual increase before FIP re-emergence, although values remained below the previously defined FIP-suggestive cutoff. Moreover, all other blood parameters remaining within reference intervals or deviations were considered not clinically relevant, highlighting the potential of AGP as an early, noninvasive monitoring parameter.

### 3.4. Optimal treatment protocol for FIP-related neurological manifestations

The optimal treatment dose and duration for cats with FIP-related neurological manifestations still need to be determined. The initial treatment protocol in this case followed a recent study showing excellent outcomes in cats with FIP-related cavitary effusions.^20^ Rapid clinical improvement and normalisation of all laboratory parameters by treatment support the initial treatment efficacy. Nevertheless, the reoccurrence of FIP-related clinical signs 17 days after initial treatment raises the question of whether a longer treatment duration, higher dose or increased frequency would have led to more sustained remission. FCoV RT-qPCR monitoring in CSF before stopping the first treatment could have detected incomplete viral clearance^6^ but was not performed due to invasiveness with respect to normalisation of neurological status.

## 4. Conclusion

This case highlights the complexity of diagnosing and treating re-emerging FIP-related neurological signs. Regular noninvasive rechecks, including AGP monitoring, allow early detection. Short-term intravenous remdesivir treatment associated with intensive care, followed by a prolonged 84-day course of high-dose oral GS-441524, led to excellent long-term outcomes despite the initially critical situation.

## Supplementary material

The following videos are available online. Video 1 D-107/42

Video 2 D-149/84

Video 3 D-149/84

## Acknowledgements

Our sincere gratitude goes to Patric Risch (the referring veterinarian) and the entire team of veterinarians and nurses from the Department of Internal Medicine and the Emergency and Intensive Care Unit at the Vetsuisse Faculty for their exceptional support and dedication throughout this case. Part of the laboratory work was performed via the logistics of the Center for Clinical Studies at the Vetsuisse Faculty of the University of Zurich.

## Author note

This case was presented in part at the 2024 ISCAID congress.

## Ethical approval

The work described in this manuscript involved the use of non-experimental (owned or unowned) animals and procedures that differed from established internationally recognised high standards (‘best practice’) of veterinary clinical care for the individual patient. The study therefore had prior ethical approval from an established (or ad hoc) committee as stated in the manuscript.

Animal study and ethical permits. The study has the following permits: 1) Approval by the governmental veterinary office (TVB No ZH124/2022; 34964, see Additional files 1 and 2). The application necessitated scientific justification, relevance to the research question, justification of the animal number, refinement techniques, definition of care and housing and compliance with all rules and regulations according to Swiss law. 2) Ethical permit (MeF-Ethik-2024-14) to legally use left over sample material sent initially for diagnostic reasons (see Additional File 3).

## Informed consent

Informed consent (verbal or written) was obtained from the owner of all animal(s) described in this work (experimental or non-experimental animals, including cadavers, tissues and samples) for all procedure(s) undertaken (prospective or retrospective studies). For any animals or people individually identifiable within this publication, informed consent (verbal or written) for their use in the publication was obtained from the people involved.

